## Supplementary figures and images for "Endogenous Retroviruses and TDP-43 Proteinopathy Form a Sustaining Feedback to Drive the Intercellular Spread of Neurodegeneration"

### Fig.S1

##### Extended Data Fig.1

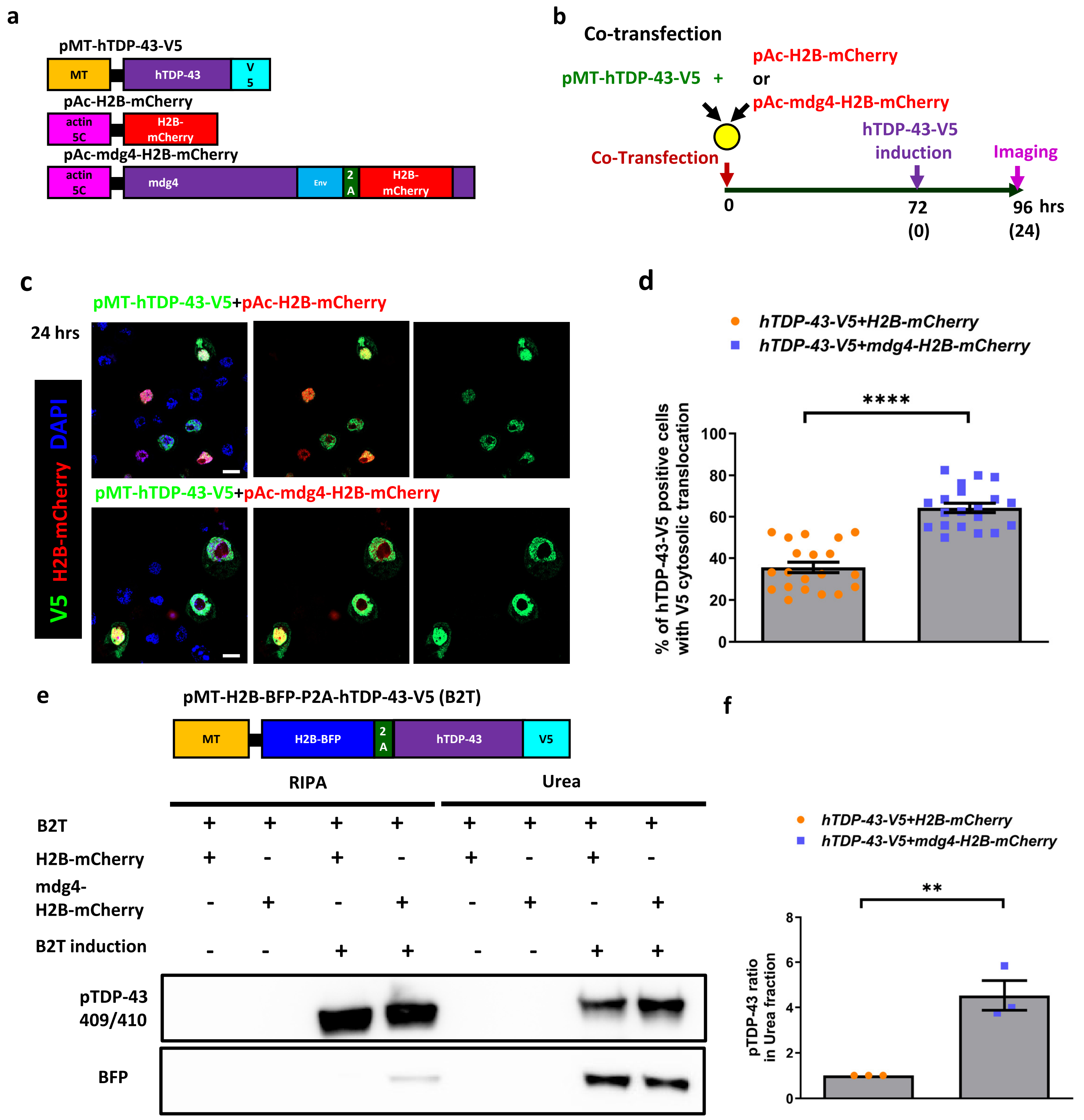

### Fig.S2

**Extended Data Fig.2**

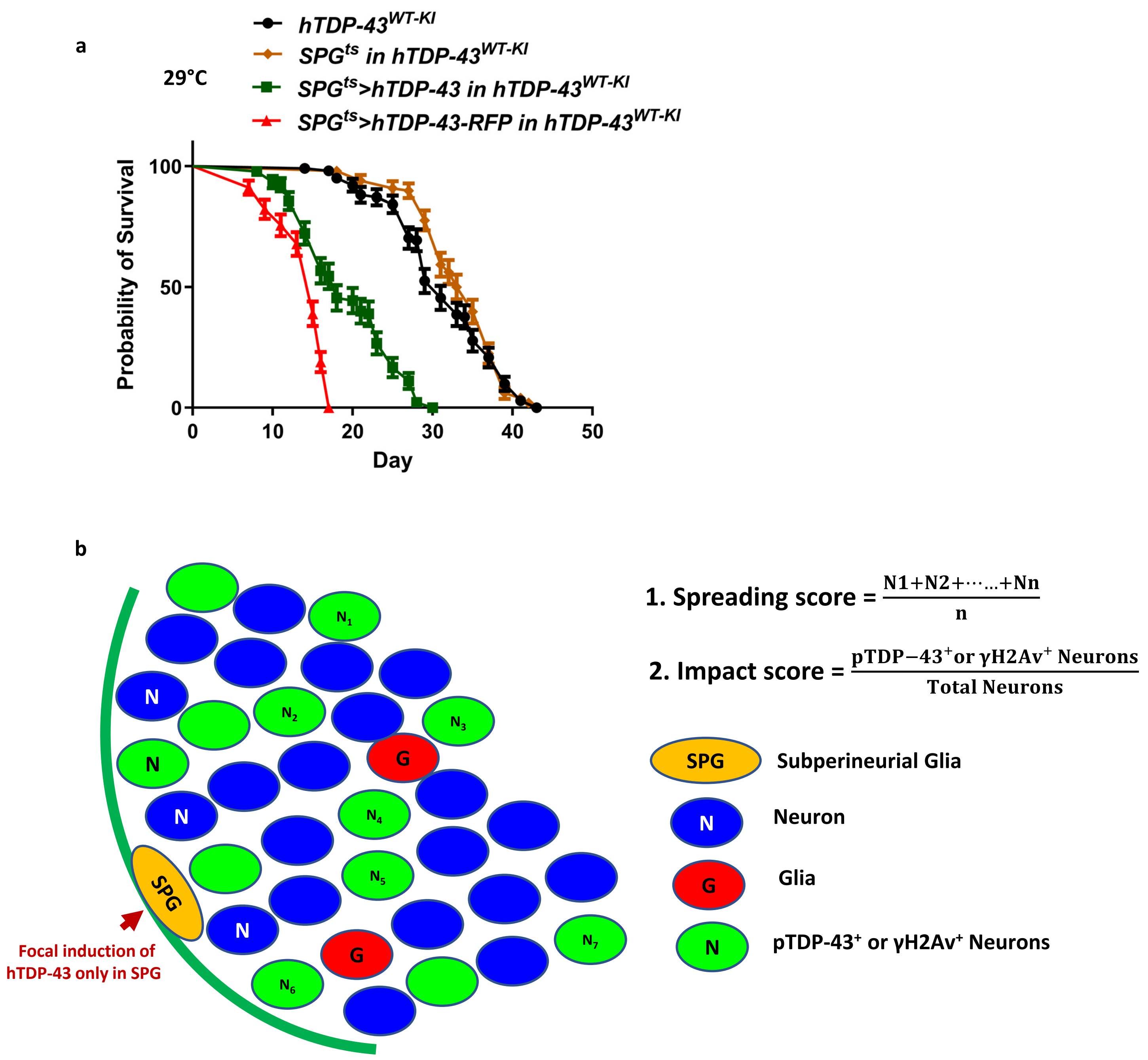

### Fig.S3

##### Extended Data Fig.3

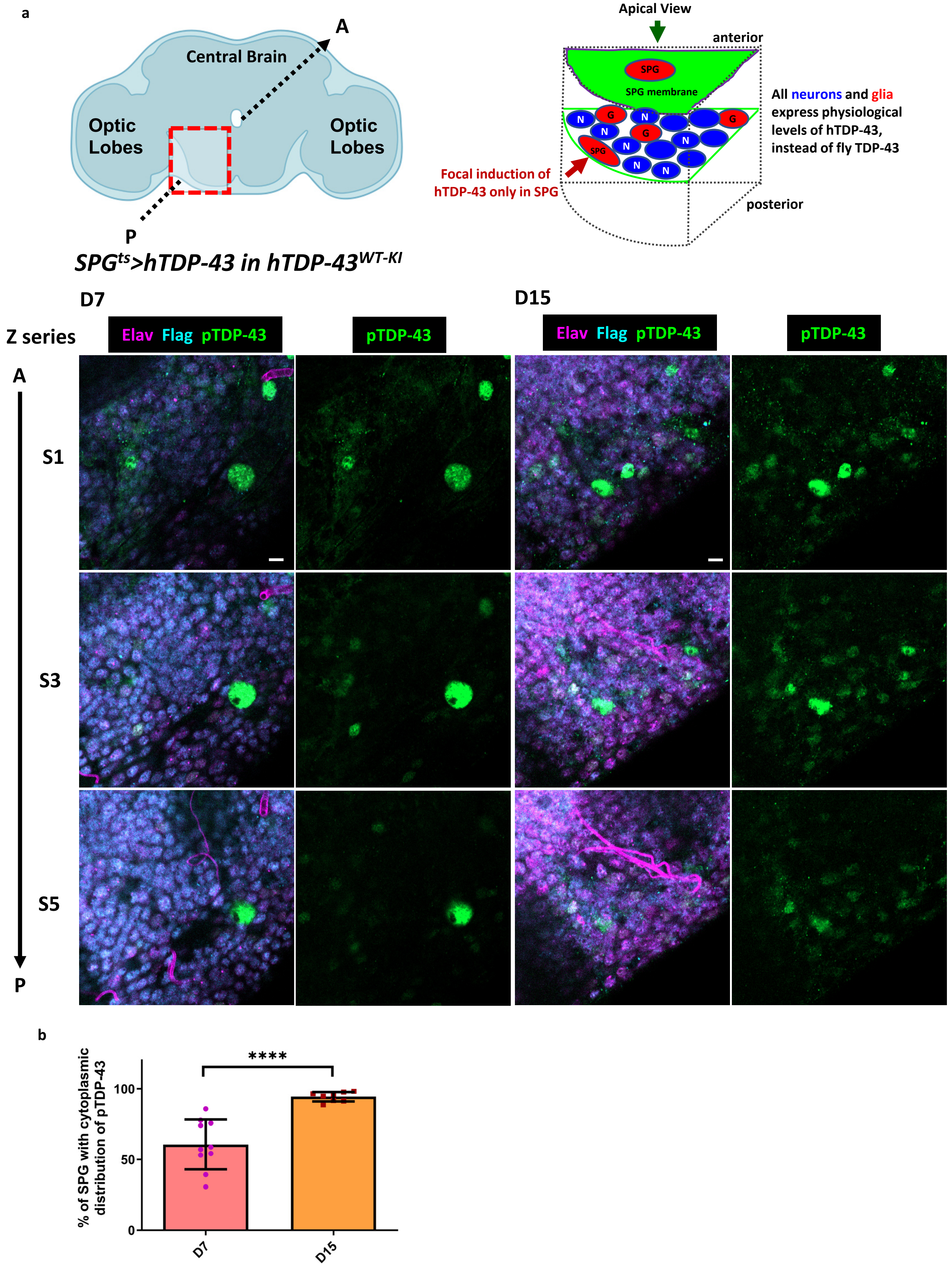

### Fig.S4

##### Extended Data Fig.4

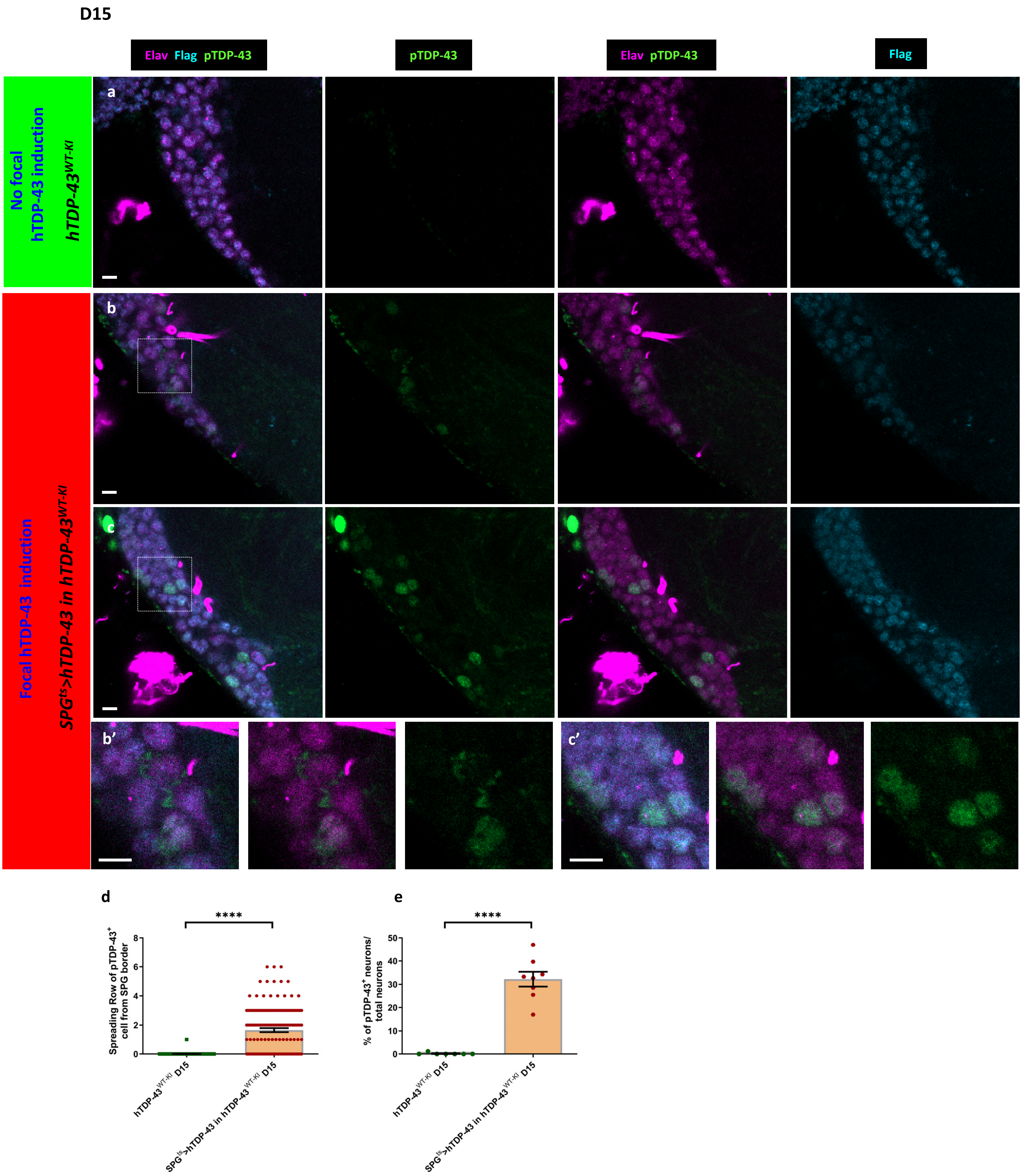

### Fig.S5

**Extended Data Fig.5**

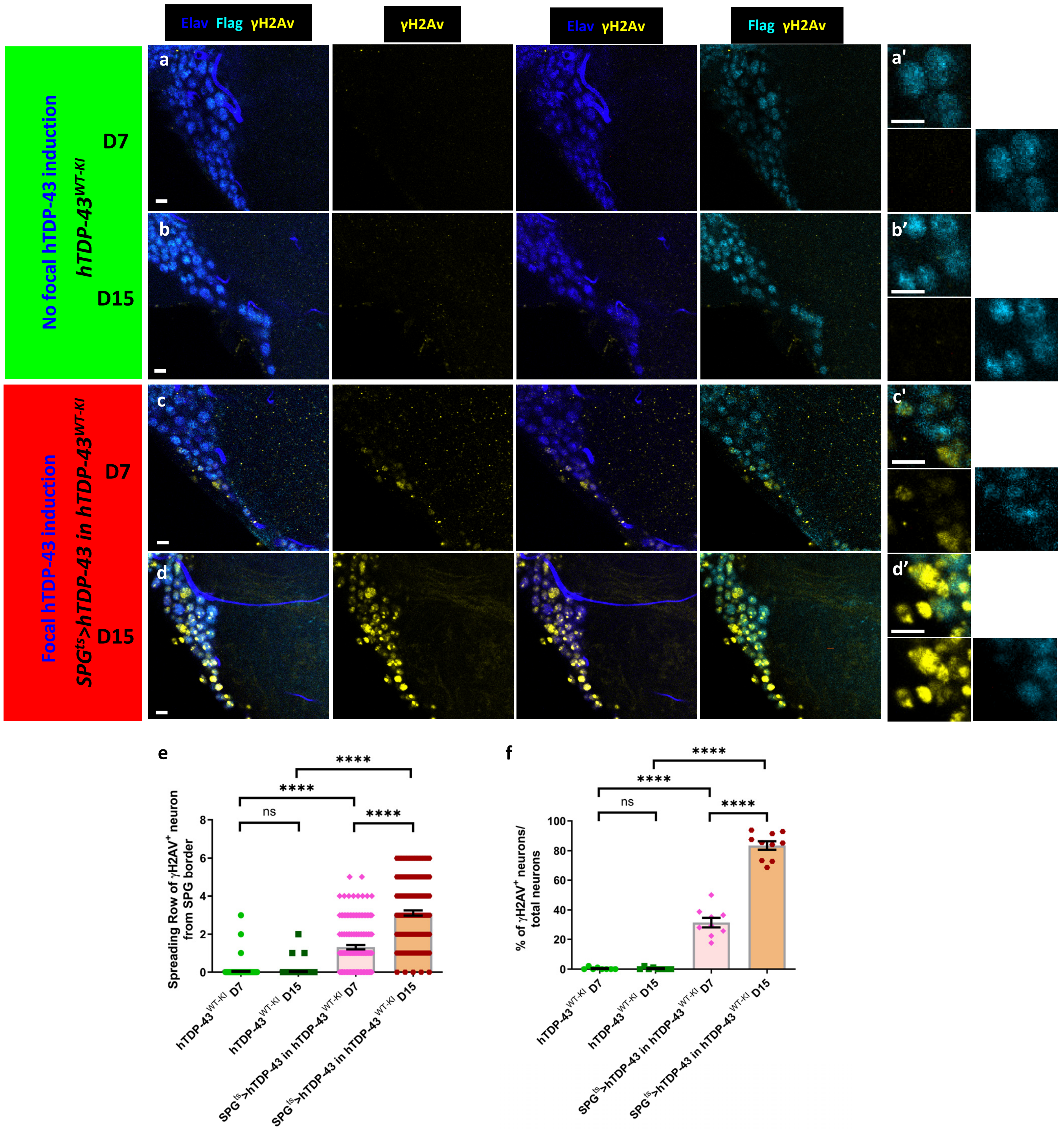

### Fig.S6

Extended Data Fig.6

a

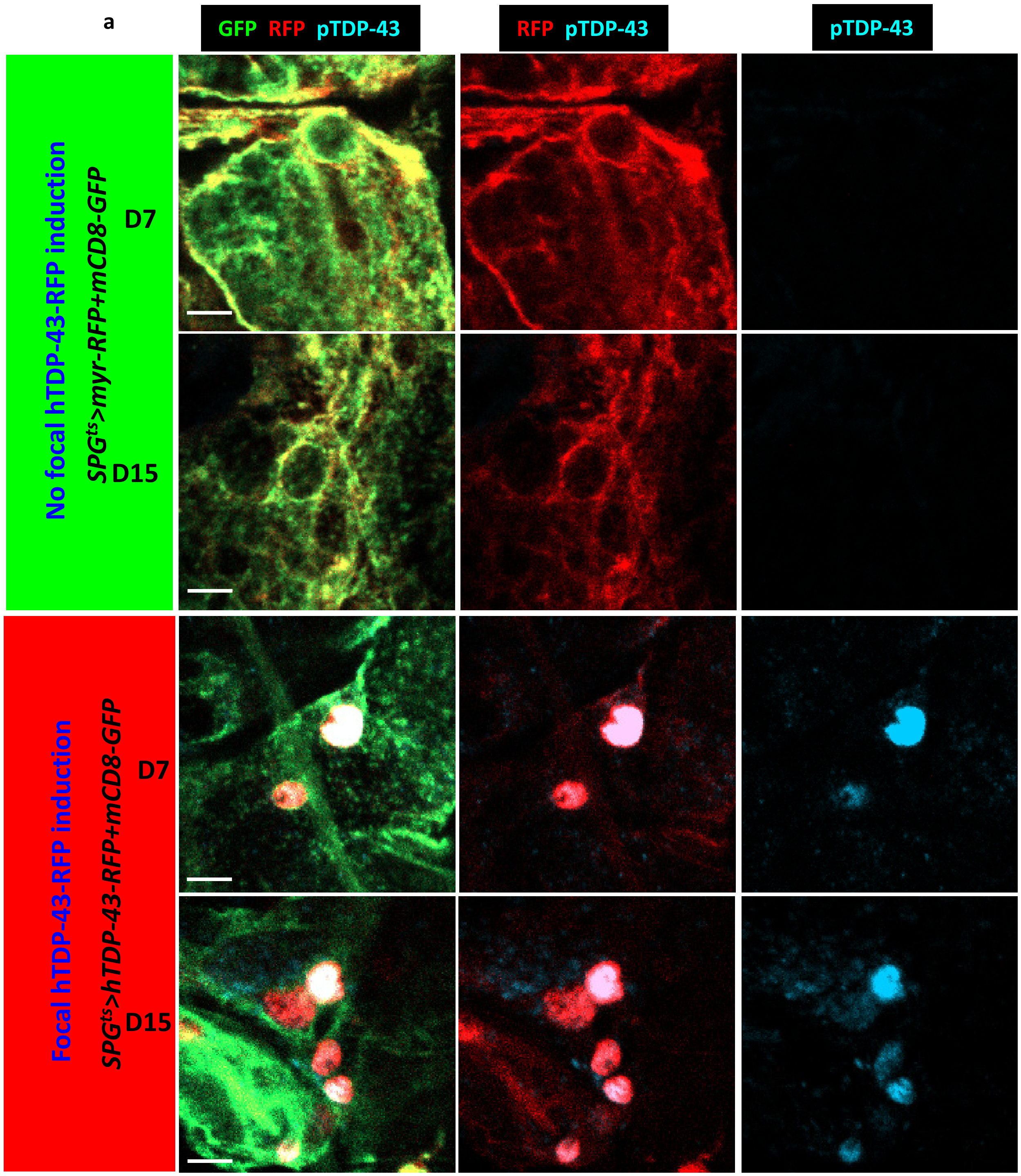


b

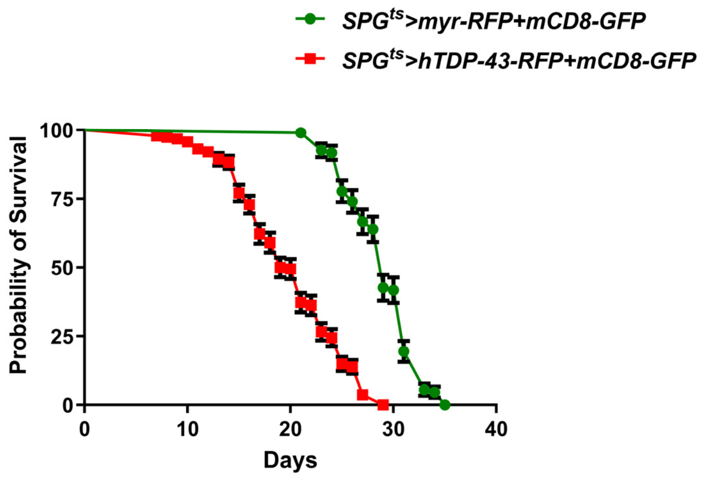

### Fig.S7

**Extended Data Fig.7**

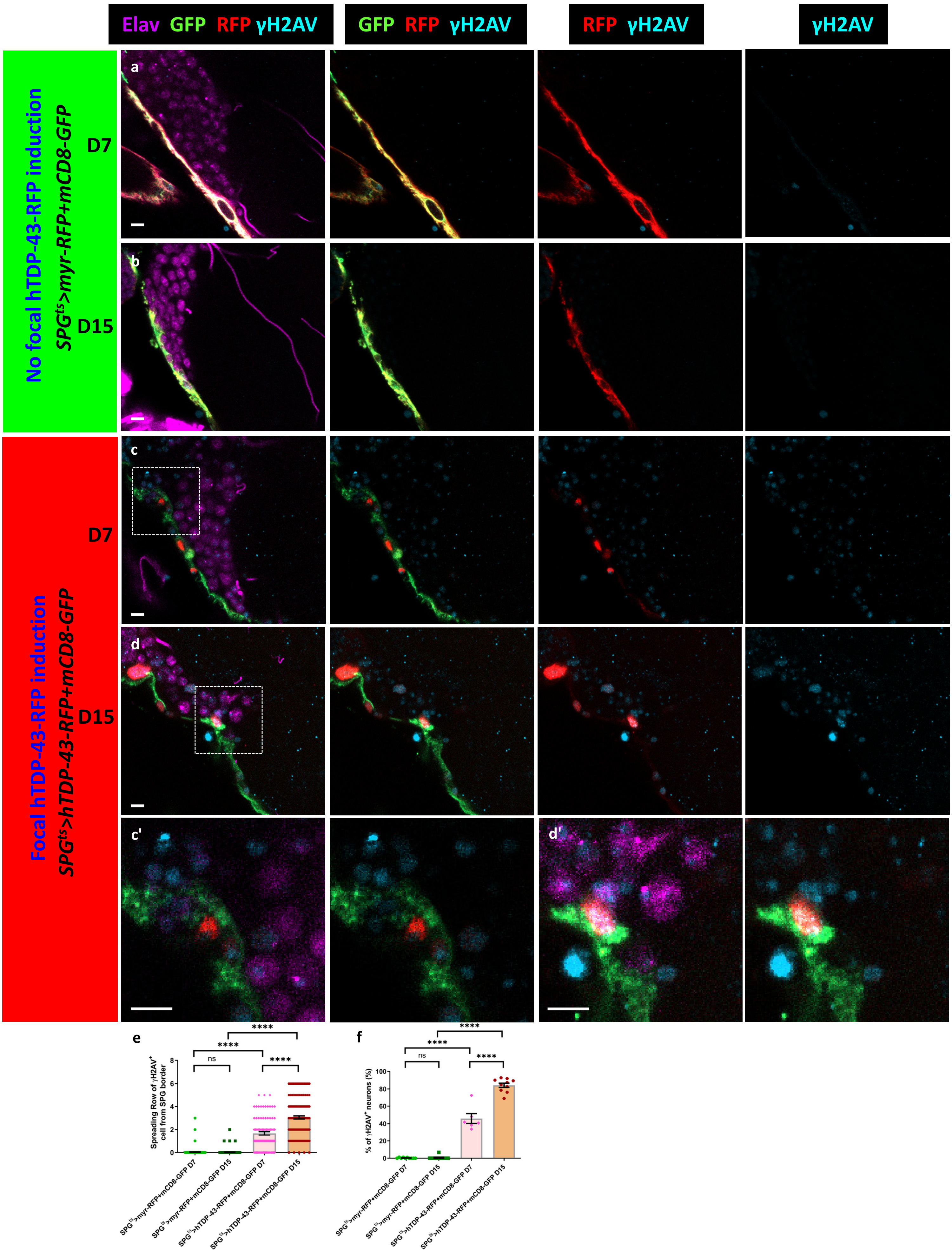

### Fig.S8

##### Extended Data Fig.8

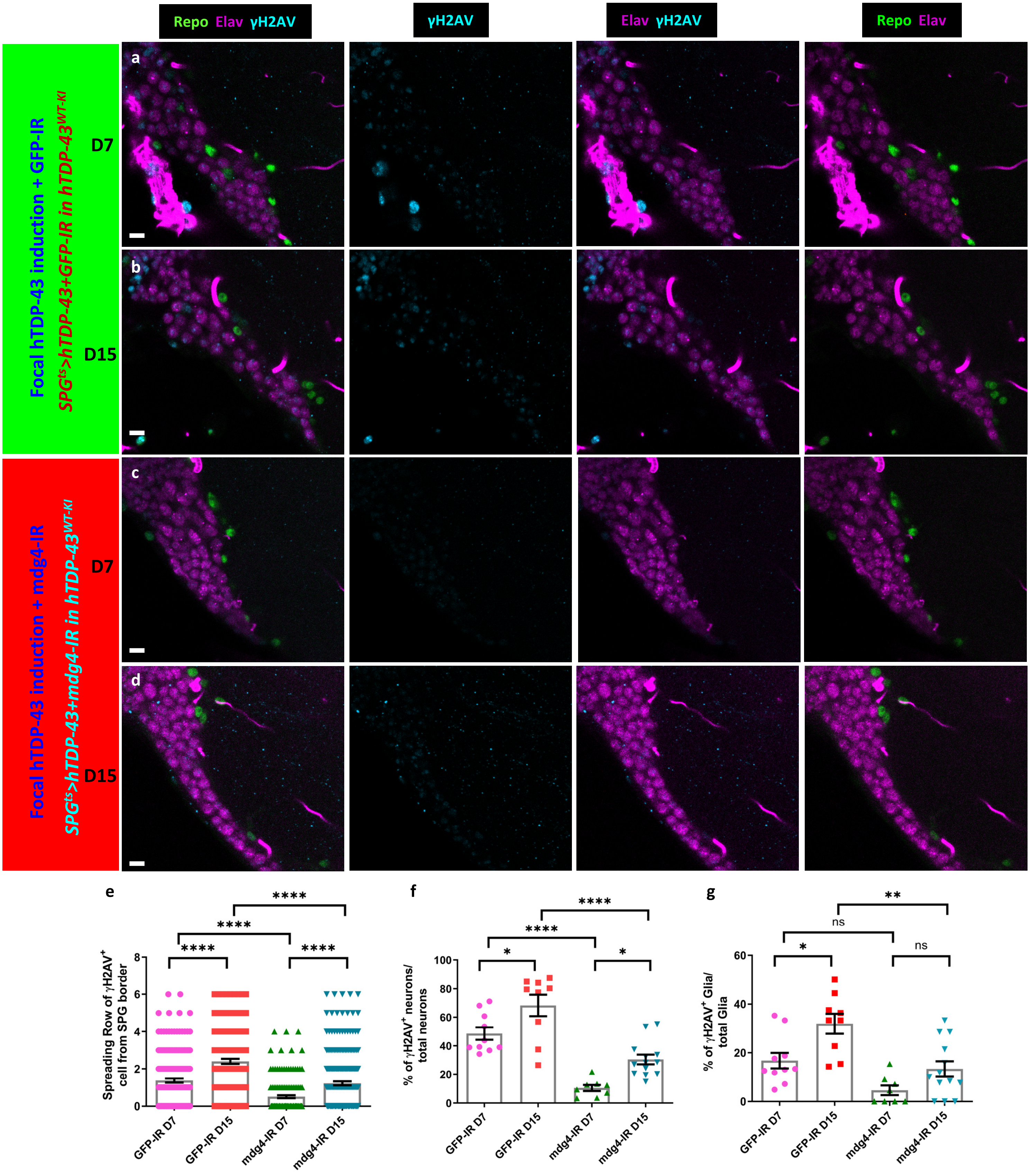

### Fig.S9

#### Extended Data Fig.9

**a**

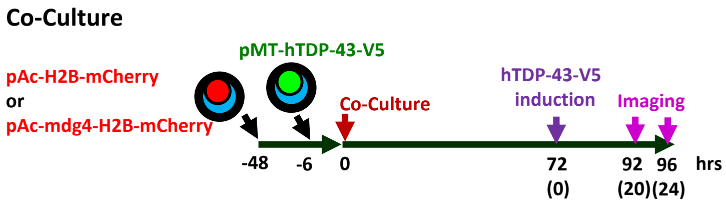


**b**

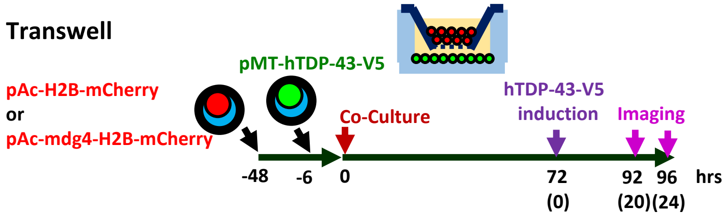
