## Supplemental Fig. Legends for "Endogenous Retroviruses and TDP-43 Proteinopathy Form a Sustaining Feedback to Drive the Intercellular Spread of Neurodegeneration"

**Extend Data Fig. 1 Expression of mdg4-ERV in *Drosophila* S2 cells causes human TDP-43 proteinopathy.** **a**, Design of an inducible hTDP-43-V5 construct is designed to co-expressed, an H2B-mCherry construct, and an mdg4-H2B-mCherry reporter. **b**, a schematic showing the timing of co-transfection and induction of mdg4-ERV or H2B-mCherry with hTDP-43-V5. **c**, Immunofluorescent images of S2 cells that were co-transfected with hTDP-43-V5 and H2B-mCherry or hTDP-43-V5 and mdg4-H2B-mCherry and stained for V5 (green), H2B-mCherry (red) and DAPI (blue). Scale bar=5µm. **d**, Quantification of cytoplasmic TDP-43-V5 translocation in S2 cells co-transfected with hTDP-43-V5+H2B-mCherry or hTDP-43-V5+mdg4-H2B-mCherry 24 hours post transfection (35.62±2.52%, n=20 examined fields; 64.31±2.23%, n=20 examined fields). Data are shown as mean ± SEM and an unpaired t-test was performed. ****p<0.01. **e**, Western blot images of RIPA-soluble and Urea-soluble fractions from S2 cells that were co-transfected with H2B-BFP-P2A-hTDP-43-V5+H2B-mCherry or H2B-BFP-P2A-hTDP-43-V5+mdg4-H2B-mCherry. pTDP-43 and BFP are imaged, with BFP serving as an internal control for a marker expressed in the same fraction of cells. **f**, Quantification of pTDP-43 levels in urea soluble fractions. The relative fold change of pTDP-43 in the urea-soluble fraction of H2B-BFP-P2A-hTDP-43-V5+mdg4-H2B-mCherry is 4.53±0.66 of H2B-BFP-P2A-hTDP-43-V5+H2B-mCherry (n=3). Data are shown as mean ± SEM and an unpaired t-test was performed. **p<0.01.

**Extend Data Fig. 2 Effects of SPG glia expression of human-TDP-43 on lifespan and schematic of spreading and impact scores to quantify spread of TDP-43 proteinopathy in the humanized *Drosophila* model.** **a**, Survival analyses for control TDP-43^WT-KI^ (n=101), SPG^ts^ (SPG-Gal4 and Gal80^ts^) in human TDP-43^WT-KI^ background (n=98), SPG^ts^>hTDP-43 in human TDP-43^WT-KI^ background (n=90) and SPG^ts^>hTDP-43-RFP in human TDP-43^WT-KI^ background (n=90). Median survival for these groups are 31, 34, 18 and 15 days respectively. Log-rank test (LR) and Gehan-Breslow-Wilcoxon (GBW) tests are used for comparisons. For human TDP-43^WT-KI^ or SPG^ts^ in human TDP-43^WT-KI^ background vs. SPG^ts^>hTDP-43 in human TDP-43^WT-KI^ background, p values are all <0.0001 in LR and GBW. For TDP-43^WT-KI^ or SPG^ts^ in human TDP-43^WT-KI^ background vs. SPG^ts^>hTDP-43-RFP in human TDP-43^WT-KI^ background, p values are all <0.0001 in LR and GBW as well. **b**, Schematic showing spatial organization of brain region where quantification were performed and the method of calculation of spreading and impact scores. Spatial locations are shown for SPG, neurons (N) and other glial subtypes (G) in the optical sections of fly adult central brain that are shown in main other figures. Also shown is the method of calculating spreading and impact scores from the optical sections of fly adult central brain. SPG, neurons (blue N), other glial subtypes (G), and pTDP-43^+^ or γH2Av^+^ neurons (green N).

**Extend Data Fig. 3 Apical view showing propagation of** **TDP-43 proteinopathy from SPG to neurons in the humanized hTDP-43 model.** **a**, Schematic of apical sections shown from the adult central brain. Immunofluorescent images from three apical sections from central brain of SPG^ts^>hTDP-43 in TDP-43^WT-KI^ background at D7 and D15 with staining for pTDP-43 (green), Flag tag on TDP-43^WT-KI^ (cyan) and neuronal cell marker Elav (violet). This view shows both the spread of pTDP-43 signal to neurons, and the appearance of cytoplasmic puncta. Scale bar=5µm. **b**, Quantification of percentage of SPG with TDP-43 proteinopathy in SPG^ts^>hTDP-43 in TDP-43^WT-KI^ background at D7 (60.69±5.57%, n=10) and D15 (94.35±1.21%, n=8). This percentage increases between D7 and D15. Data are shown as mean ± SEM and an unpaired t-test is performed. ****p<0.01.

**Extend Data Fig. 4 Propagation of TDP-43 proteinopathy in the humanized *Drosophila* hTDP-43 model confirmed with an alternate pTDP-43 antibody.** **a-c**, Immunofluorescent images from central brain sections with staining for pTDP-43 (green), The Flag tag on TDP-43^WT-KI^ (cyan) and the neuronal cell marker Elav (violet). Scale bar=5µm. **a**, TDP-43^WT-KI^ at D15. **b-c**, SPG^ts^>hTDP-43 in a TDP-43^WT-KI^ background at D15. **d**, Quantification of pTDP-43 spreading scores of TDP-43^WT-KI^ and SPG^ts^>hTDP-43 in TDP-43^WT-KI^ background at D15 (0.01±0.01, n=110; 1.65±0.14, n=136). Data are shown as mean ± SEM and an unpaired t-test was performed. ****p<0.01. **e**, Quantification of pTDP-43 impact scores of TDP-43^WT-KI^ and SPG^ts^>hTDP-43 in TDP-43^WT-KI^ background at D15 (0.16±0.16%, n=7; 32.25±3.19%, n=8). Data are shown as mean ± SEM and an unpaired t-test was performed. ****p<0.01.

**Extend Data Fig. 5 Focal induction of TDP-43 proteinopathy in SPG induces DNA damage in neurons.** **a-d’**, Immunofluorescent images from central brain sections with staining for γH2Av (yellow), Flag tag on TDP-43^WT-KI^ (cyan) and neuronal cell marker Elav (blue). Scale bar=5µm. **a-b’**, TDP-43^WT-KI^ at D7 and D15. **c-d’**, SPG^ts^>hTDP-43 in TDP-43^WT-KI^ background at D7 and D15. **a’-b’**, Enlarged images from selected regions of a and b. **c’-d’**, Enlarged images from the selected region of c and d. **e**, Quantification of γH2Av spreading scores of TDP-43^WT-KI^ and SPG^ts^>hTDP-43 in TDP-43^WT-KI^ background at D7 (0.04±0.03, n=139; 1.31±0.11, n=140) and D15 (0.03±0.02, n=134; 3.11±0.14, n=144). Data are shown as mean ± SEM and a two-way ANOVA with multiple comparisons was performed. ****p<0.0001; ns, no significant difference. **f**, Quantification of γH2Av impact scores of TDP-43^WT-KI^ and SPG^ts^>hTDP-43 in TDP-43^WT-KI^ background at D7 (0.42±0.29%, n=8; 31.56±3.24%, n=9) and D15 (0.42±0.29%, n=8; 83.54±2.83%, n=10). Data are shown as mean ± SEM and a two-way ANOVA with multiple comparisons was performed. ****p<0.0001; ns, no significant difference.

**Extend Data Fig. 6 Focal induction of hTDP-43-RFP in SPG causes lifespan reduction and cell autonomous hTDP-43 proteinopathy in SPG.** **a**, Apical view showing Immunofluorescent images from central brain sections of SPG^ts^>myr-RFP+mCD8-GFP and SPG^ts^>hTDP-43-RFP+mCD8-GFP with staining for pTDP-43 (cyan), RFP (red) and membrane GFP (green) at D7 and D15. Scale bar=5µm. **b**, Survival assays for SPG^ts^>myr-RFP+mCD8-GFP (n=108) and SPG^ts^>hTDP-43-RFP+mCD8-GFP (n=188) exhibit a median survival of 29 and 19.5 days respectively. Log-rank test (LR) and Gehan-Breslow-Wilcoxon (GBW) test are used. For SPG^ts^>myr-RFP+mCD8-GFP vs SPG^ts^>hTDP-43-RFP+mCD8-GFP, p values are all <0.0001 in LR and GBW.

**Extend Data Fig. 7 Induction of hTDP-43-RFP in SPG causes nonautonomous spread of DNA damage to neurons.** **a-d**, Immunofluorescent images from of central brain sections of SPG^ts^>myr-RFP+mCD8-GFP and SPG^ts^>hTDP-43-RFP+mCD8-GFP with staining for γH2Av (cyan), RFP (red), membrane GFP (green) and neuronal cell marker Elav (violet) at D7 and D15. Scale bar=5µm. **a-b**, SPG^ts^>myr-RFP+mCD8-GFP at D7 and D15. **c-d**, SPG^ts^>hTDP-43-RFP+mCD8-GFP at D7 and D15. **c’-d’**, Enlarged images from the indicated regions of c and d. **e**, Quantification of γH2Av spreading scores of SPG^ts^>myr-RFP+mCD8-GFP and SPG^ts^>hTDP-43-RFP+mCD8-GFP at D7 (0.04±0.02, n=164; 1.67±0.15, n=88) and D15 (0.03±0.01, n=180; 3.06±0.13, n=139). Data are shown as mean ± SEM and a two-way ANOVA with multiple comparisons was performed. ****p<0.0001; ns, no significant difference. **f**, Quantification of γH2Av impact scores of SPG^ts^>myr-RFP+mCD8-GFP and SPG^ts^>hTDP-43-RFP+mCD8-GFP at D7 (0.33±0.17%, n=10; 45.88±5.63%, n=6) and D15 (0.62±0.62%, n=11; 84.22±2.44%, n=10). Data are shown as mean ± SEM and a two-way ANOVA with multiple comparisons was performed. ****p<0.0001; ns, no significant difference.

**Extend Data Fig. 8 Mdg4-ERV expression in SPG is required for the propagation of DNA damage from SPG to neurons.** **a-d**, Immunofluorescent images from central brain sections with staining for γH2Av (cyan), neuronal cell marker Elav (violet) and glial cell marker Repo (green). Scale bar=5µm. **a-b**, SPG^ts^>hTDP-43+GFP-IR in human TDP-43^WT-KI^ background at D7 (a) and D15 (b). **c-d**, SPG^ts^>hTDP-43+mdg4-IR in human TDP-43^WT-KI^ background at D7 (c) and D15 (d). **e**, Quantification of γH2Av spreading scores of SPG^ts^>hTDP-43+GFP-IR in human TDP-43^WT-KI^ background and SPG^ts^>hTDP-43+mdg4-IR in human TDP-43^WT-KI^ background at D7 (1.37±0.11, n=179; 0.50±0.07, n=146) and D15 (2.41±0.14, n=148; 1.22±0.10, n=231). Data are shown as mean ± SEM and a two-way ANOVA with multiple comparisons was performed. ****p<0.0001. **f**, Quantification of γH2Av impact scores of SPG^ts^>hTDP-43+GFP-IR in human TDP-43^WT-KI^ background and SPG^ts^>hTDP-43+mdg4-IR in human TDP-43^WT-KI^ background at D7 (48.61±4.35%, n=10; 10.51±2.14%, n=8) and D15 (68.33±7.54%, n=9; 30.43±3.42%, n=13). Data are shown as mean ± SEM and a two-way ANOVA with multiple comparisons was performed. *p<0.05; ****p<0.0001. **g**, Quantification of γH2Av impact scores on other glial cells of SPG^ts^>hTDP-43+GFP-IR in TDP-43^WT-KI^ background and SPG^ts^>hTDP-43+mdg4-IR in human TDP-43^WT-KI^ background at D7 (16.76±3.21%, n=10; 4.52±2.01%, n=8) and D15 (31.93±4.10%, n=9; 13.32±3.17%, n=13). Data are shown as mean ± SEM and a two-way ANOVA with multiple comparisons was performed. *p<0.05; **p<0.01; ns, no significant difference.

**Extend Data Fig. 9 Experimental schematic of co-culture and transwell assays of mdg4-ERV viral transmission to induce human** **TDP-43 proteinopathy in recipient cells.** **a**, H2B-mCherry or mdg4-H2B-mCherry were transfected in the producing cells and then co-cultured with hTDP-43-V5 recipient cells. **b**, H2B-mCherry or mdg4-H2B-mCherry were transfected in the producing cells and then cultured in opposition to hTDP-43-V5 recipient cells in a transwell system.
