## Supplement with Methods for "Endogenous Retroviruses and TDP-43 Proteinopathy Form a Sustaining Feedback to Drive the Intercellular Spread of Neurodegeneration"

**Fly strains and husbandry**

We obtained SPG-Gal4 (R54C07-Gal4, #50472) and UAS-myr-RFP (#7119) from Bloomington Drosophila Stock Center. The hTDP-43-KI^1^ and UAS-hTDP-43-RFP^2^ were generous gifts from Professor David Morton (Oregon Health & Science University) and Professor Jane Y. Wu (Northwestern University School of Medicine). The following stocks, UAS-GFP-IR, UAS-gypsy-IR, UAS-hTDP-43, tub-Gal80^ts^ and UAS-mCD8-GFP were used in our previous studies^3-6^. To prevent artifacts from genetic variation between groups, all strains used in this study were backcrossed to our laboratory wild-type strain, Canton-S derivative w^1118^ (*isoCJ1*), for at least five generations. Male flies were chosen as the experimental subjects throughout the study. All flies for TARGET^7^ temperature-shift experiments (*SPG-Gal4* combined with *tub-Gal80^ts^*) were raised in a 21°C incubator from the embryonic stage^7^. All flies for TARGET experiments were immediately shifted to the permissive temperature at 29°C after eclosion, which was designated as day 0. These TARGET flies were then incubated at 29°C for lifespan assays or dissected at desired time points for analyzing the patterns of molecular markers dependent on the experimental designs.

**Drosophila S2 expression constructs**

To generate the inducible hTDP-43-V5 construct, the full length of hTDP-43 was PCR amplified but the stop codon was removed in order to generate an in frame fusion protein with the V5 tag provided by the pMT-V5-6xHis plasmid (Invitrogen, V412020). To generate the pMT-H2B-BFP-P2A-hTDP-43-V5 construct, the H2B and BFP were separately PCR amplified from pAc-H2B-mCherry^8^ and moxBFP (Addgene, 68064) constructs and then fused by PCR to generate the H2B-BFP fragment. The P2A was separately added to the C-terminal of H2B-BFP and N-terminal of hTDP-43 by PCR. The final H2B-BFP-P2A-hTDP-43 was then created by PCR fusing H2B-BFP-P2A and P2A-hTDP-43 fragments and then cloned into the inducible pMT-V5-6xHis plasmid. The pAc-H2B-mCherry-HA and pAc-gypsy-H2B-mCherry-HA constructs were generated described previously^8^.

**Mammalian SH-SY5Y expression constructs**

To generate the constitutively expressing pcDNA3.1-H2B-mCherry-HA, the H2B-mCherry-HA was moved from pAc-H2B-mCherry-HA by restriction enzyme NotI and reinserted the H2B-mCherry-HA DNA fragment into the NotI site of pcDNA3.1 (Thermo Fisher Scientific, V79020). The backbone of HERV-K (pcDNA3.1-HERV-K) was obtained from Professor Nath’s laboratory^9^ and used as the template to generate the constitutive pcDNA3.1-HERV-K-P2A-H2B-mCherry-HA plasmid. In short, the P2A-H2B-mCherry-HA fragment was PCR amplified and integrated downstream of Env of pcDNA3.1-HERV-K while removing stop codon of Env.

**Adult fly survival assays**

Adult male flies with desired genotypes raised at 21°C were collected immediately after eclosion and incubated in a 29°C incubator for survival assay as addressed in previous studies^3, 4^. In short, ten adult flies of a given genotype were housed within one vial, and more than 100 flies in total for experimental group were used for each experiment. The surviving flies from each vial were flipped into fresh vials with fly food every other day, and dead flies at specific time points were recorded for the final survival curve analysis. The Log-Rank (Mantel-Cox) test and the Gehan-Breslow-Wilcoxon test were used to compare the significance of survival curves.

**Immunostaining of adult fly brains**

Adult brains at specific time points for each experiment were dissected, and immunofluorescent staining was performed as previously described^3^. Adult brains were dissected in ice-cold phosphate-buffered-saline (PBS) and then transferred into 4% paraformaldehyde (Electron Microscopy Sciences #15713) PBS solution (1XPBS with 4% paraformaldehyde and 0.2% Triton-X-100 (SIGMA-ALDRICH)) and incubated for 30 minutes under vacuum twice. After fixation, dissected brains were washed three times for 10 minutes with 1XPBST wash solution (1XPBS with 1% Triton-X-100 and 3% NaCl). Brain samples were then incubated in blocking solution (1XPBST with 10% normal horse serum) overnight at 4°C on a nutator. After blocking, dissected brains were transferred into primary antibodies and incubated overnight at 4°C with the following dilutions: mouse anti-Repo (1:10, Developmental Studies Hybridoma Bank 8D12), rat anti-Elav (1:10, Developmental Studies Hybridoma Bank 7E8A10), mouse anti-Elav (1:10, Developmental Studies Hybridoma Bank 9F8A9), mouse anti-Flag (1:100, SIGMA-ALDRICH F3165), rabbit anti-pTDP-43 (1:100, SIGMA-ALDRICH SAB4200223), rabbit anti-pTDP-43 (1:100, proteintech 22309-1-AP) and rabbit anti-γH2Av (1:150, Rockland Immunochemicals 600-401-914) in 1XPBST washing solution with 10% normal horse serum. Samples were then washed with 1XPBST four times, 15 minutes each. Fly brains were incubated in secondary antibody solution (1XPBST washing solution with 10% normal horse serum) at 4°C overnight. After washing with 1XPBST for 4X 15 minutes, fly brains were mounted in FocusClear (CelExplorer), imaged using a Zeiss LSM 800 with Airyscan mode, and acquired images were processed by the Zeiss Zen software package.

**Quantification of spreading and severity of TDP-43 pathology and DNA damage in the adult fly brain**

To standardize the brain region for comparing changes of molecular markers over time and between different genetic groups, patterns of molecular markers in a defined 5595.7825µm^2^ (77.99µm x 71.75µm) area (as shown in Fig.1b, one corner of the selected rectangle was chosen at the overlap between the central brain and optic lobe) of each brain were quantified. In order to monitor the spread of pTDP-43 or γH2Av, the farthest cell within the same row relative to the SPG membrane margin with positive molecular markers was given a number dependent on its sequential cell position away from the SPG membrane within its row (as shown in the Fig. 1i). All numbers assigned to each row in the selected brain region at a particular time window were averaged to calculate a “spreading score” of that genotype and time-point. The “impact score” was designed to compare the fraction of cells labeled with a given marker in a selected genotype and time point. In short, the neuronal cells with pTDP-43 or DNA damage markers within the selected brain region were counted and divided by the total number of Elav+ neurons in that region to calculate a percentage of neuronal cells labelled.

**Cell culture and transfection conditions**

*Drosophila* S2 cells (Thermo Fisher Scientific, R69007) were maintained in Schneider’s Drosophila medium (Thermo Fisher Scientific, 21720001) supplemented with 10% fetal bovine serum (FBS) (Thermo Fisher Scientific, 10438026) and 100U/ml Penicillin-Streptomycin (Thermo Fisher Scientific, 15140122) in a 25°C incubator. Human neuroblastoma cells, SH-SY5Y (ATCC, CRL-2266), were grown in Dulbecco’s Modified Eagle Medium/Nutrient Mixture F-12 (DMEM/F-12) (Thermo Fisher Scientific, 11320033) with 10% FBS and 100U/ml Penicillin-Streptomycin in a 37°C incubator with 5% CO_2_ supplementation.

Conditions for S2 cells transfection were modified from our previous studies^6, 8^. 5x10^5^ cells were seeded onto coverslips coated with 0.5mg/ml concanavalin A (ConA) and placed in the 6 well culture plate overnight prior to the transfection. Single or co-transfections with desired construct combinations (pAc-H2B-mCherry, pAc-gypsy-P2A-H2B-mCherry and pMT-hTDP-43-V5) were performed with 1µg of DNA each and Effectene transfection reagents (Qiagen, 301427). Post 24 hours transfection, transfection complexes were washed away and supplied with the fresh complete medium containing 700µM CuSO_4_ to induce hTDP-43-V5 expression.

Transfection conditions of SH-SY5Y were as follows: 8x10^5^ cells were seeded onto coverslips coated with 0.1mg/ml Poly-D-Lysine and placed in 6 well culture plates overnight prior to transfection. Single transfection was performed under the manufacturer’s guideline with a mixture of 4µg pcDNA3.1-H2B-mCherry or pcDNA3.1-HERV-K-P2A-H2B-mCherry with 12µl TransIT-2020 reagent in a 1:3 ratio (Mirus, MIR 5400). After 24 hours of incubation, transfection complexes were washed away, and fresh complete medium was supplied daily until the desired time points for analysis.

**Co-culture and transwell setup for cellular assays of Drosophila S2 cells**

In order to expand the donor cell population and enhance the production of virus-like particles (VLPs), S2 cells were transfected with pAc-H2B-mCherry or pAc-gypsy-P2A-H2B-mCherry 48 hours before co-culture or transwell assays were performed (as flowchart shown in Extended Data Fig.9). For the hTDP-43-V5 recipient population, the inducible pMT-hTDP-43-V5 was delivered with a 6 hour short transfection prior to the experimental setup. For co-culture, 5x10^5^ of donor (pAc-H2B-mCherry or pAc-gypsy-P2A-H2B-mCherry) and recipient (pMT-hTDP-43-V5) cells were seeded at a 1:1 ratio onto the ConA-coated coverslip and maintained for 72 hours to enhance the entrance of VLPs into recipient cells. At 20 or 24 hours prior to the experimental time, the culture medium was replaced with a medium containing 700µM CuSO_4_ to induce hTDP-43-V5 expression in recipient cells. For transwell assays, 5x10^5^ of recipient (pMT-hTDP-43-V5) cells were plated onto ConA-coated coverslips and placed in the bottom well. At the same time, 1x10^6^ of donor (pAc-H2B-mCherry or pAc-gypsy-P2A-H2B-mCherry) cells were placed into the upper well. 20 or 24 hours prior to analysis, CuSO_4_ was added to induce hTDP-43-V5 expression in recipient cells.

**Immunostaining of Drosophila S2 and human SH-SY5Y cells**

S2 or SH-SY5Y cells on the ConA or Poly-D-Lysine coated coverslips were washed three times with 1XPBS and immediately fixed in 4% paraformaldehyde (Electron Microscopy Sciences #15713) PBS solution (1XPBS with 4% paraformaldehyde and 0.2% Triton-X-100 (SIGMA-ALDRICH)) for 10 minutes on a nutator. After fixation, cells were washed three times with 1XPBST wash solution (1XPBS with 1% Triton-X-100 and 3% NaCl) and then incubated for 1 hour in blocking solution (1XPBST with 10% normal horse serum) at room temperature. Cells were transferred into primary antibodies in 1XPBST washing solution with 10% normal horse serum after blocking and incubated for 1 hour at room temperature with the following dilutions: mouse anti-V5 (1:250, Thermo Fisher Scientific R960-25), rabbit anti-TDP-43 (1:200, proteintech 10782-2-AP) and rabbit anti-pTDP-43 (1:200, SIGMA-ALDRICH SAB4200223), rabbit anti-pTDP-43 (1:500, proteintech 22309-1-AP) and mouse anti-HERV-K-Env (1:500, AUSTRAL Biologicals). Samples were then washed with 1XPBST three times for 10 minutes each. Cells were incubated within secondary antibody solution (1XPBST washing solution with 10% normal horse serum) with DAPI at room temperature for 1 hour. After washing three times with 1XPBST for 10 minutes each, cells were mounted in ProLong Diamond Antifade Mountant (Thermo Fisher Scientific), imaged using a Zeiss LSM 800 with Airyscan mode, and acquired images were processed by Zeiss Zen software package.

**Sequential fractionation of cellular TDP-43 aggregates for Western Blot analysis**

The soluble and insoluble TDP-43 fractions within the S2 cells were sequentially extracted dependent on the solubilities in the extraction buffers as previously descripted^10^. Transfected S2 cells were washed, collected with 1XPBS, and then homogenized in ice-cold RIPA buffer (Thermo Fisher Scientific, 89900) with protease inhibitors (Thermo Fisher Scientific, A32965) and phosphatase inhibitors (Roche, 4906845001). Cell lysates were centrifuged at 16000g for 30 minutes at 4°C, and the supernatant was saved as the RIPA-soluble fraction. The RIPA-insoluble pellets were washed with RIPA buffer by centrifuging at 16000g for 15 minutes at 4°C. Supernatants were discarded and the pellets were extracted and homogenized with urea buffer (7M Urea, 4% CHAPS, 2M Thiourea, 30mM Tris, pH=8.5). The lysates were centrifuged at 16000g for 30 minutes at 4°C. The supernatant was kept as the RIPA-insoluble and urea-soluble fraction.

30µg protein samples from each cellular fraction were separated by 10% polyacrylamide gel (BIO-RAD, 4561034) and followed by a canonical Western Blot protocol. In short, membranes were blocked in 1XTris Buffered Saline-Tween-20 (TBST) (1XTBS+0.2% Tween-20) with 5% SlimFast (chocolate flavor) at room temperature for 30 minutes. After blocking, membranes were incubated with primary antibodies mouse anti-GFP (1:1000, Thermo Fisher Scientific MA5-15256), and rabbit anti-pTDP-43 (1:1000, proteintech 22309-1-AP)) in 5% SlimFast -TBST for 1 hour at room temperature or overnight at 4°C. After washing three times with 1XTBST, membranes were incubated with secondary antibodies (goat anti-mouse-HRP (Jackson ImmunoResearch Laboratories, 115-035-174) or goat anti-rabbit-HRP (Jackson ImmunoResearch Laboratories, 111-035-144) at 1:1000 or 1:2000 dilutions) for 1 hour at room temperature. Signals were developed in chemiluminescent HRP substrate (Millipore-Sigma, WBKLS0100) and imaged with Sapphire Biomolecular Imager (Azure Biosystems).

**Statistics and reproducibility**

Statistical analyses for different experimental setups were performed using GraphPad Prism 9. Specific statistical analyses used for comparing the significance between experimental groups were addressed in the corresponding figure legends and the experimental reproducibility was also listed in the figure legends. The significant levels were indicated as star numbers in the following format: *p<0.05, **p<0.01, ****p<0.001 and ****p<0.0001.

**Data availability**

This study did not generate/analyze large numerical datasets and code.

**Acknowledgements**

We thank Professor Avindra Nath and Dr. Wenxiu Li (National Institutes of Health) for providing the HERV-K backbone. We also thank Professor David Morton (Oregon Health & Science University) and Professor Jane Y. Wu (Northwestern University School of Medicine) sending us the hTDP-43^WT-KI^ and UAS-hTDP-43-RFP transgenic strains. We thank Roger Sher, Wanhe Li, Grigori Enikolopov, Maria de la Paz Fernandez and Tim Mosca for comments on the manuscript, and Enas Gad Elkarim, Lillian Talbot, Sarah Krupp, Meng-Fu Shih and Jorge Aspurua for helpful discussions. This work was supported by NIA awards RF1AG057338 and RF1AG076493 to J.D.

**Author contributions**

Y.H.C. and J.D. designed the experiments. Y.H.C. performed all of the experiments and collected and analyzed all the data. Y.H.C. and J.D. wrote the manuscript.

**Competing interests**

The authors declare no competing interests.
